# Osteoclasts contribute to early development of chronic inflammation by promoting dysregulated hematopoiesis and myeloid skewing

**DOI:** 10.1101/2020.12.09.418137

**Authors:** Maria-Bernadette Madel, Lidia Ibáñez, Thomas Ciucci, Julia Halper, Majlinda Topi, Henri-Jean Garchon, Matthieu Rouleau, Christopher G Mueller, Laurent Peyrin-Biroulet, David Moulin, Claudine Blin-Wakkach, Abdelilah Wakkach

**Affiliations:** Université Côte d’Azur, CNRS, LP2M, Nice, France; David H. Smith Center for Vaccine Biology and Immunology, Department of Microbiology and Immunology, University of Rochester Medical Center, Rochester, NY 14642, USA; Université Paris-Saclay, UVSQ, INSERM, Infection et inflammation, 78180, Montigny-Le-Bretonneux, France; CNRS UPR 3572, IBMC, University of Strasbourg, 67000 Strasbourg, France; Service d’hépato-gastroentérologie, CHRU de Nancy, Vandœuvre Les Nancy, France; Université de Lorraine, INSERM, Vandœuvre Les Nancy, France; Université de Lorraine, CNRS, IMoPA, Université de Lorraine, Vandœuvre Les Nancy, France

## Abstract

Increased myelopoiesis is a hallmark of many chronic inflammatory diseases. However, the mechanisms involved in the myeloid skewing of hematopoiesis upon inflammation are still incompletely understood. Here, we identify an unexpected role of bone-resorbing osteoclasts in promoting hematopoietic stem cell (HSC) proliferation and differentiation towards myeloipoiesis in the early phases of chronic colitis. RNAseq analysis revealed that osteoclasts in colitis differ from control ones and overexpress genes involved in the remodeling of HSC niches. We showed that colitic osteoclasts modulate the interaction of HSCs with their niche and promote myeloid differentiation. Increased osteoclast activity was correlated with an augmentation of myelopoiesis in patients with chronic colitis. Therapeutic blockade of osteoclasts reduced HSC proliferation and myeloid skewing and resulted in a decreased inflammation and severity of colitis. Together, these data identify osteoclasts as potent regulators of HSCs and promising target in chronic colitis.

## Introduction

Several recent studies have highlighted the ability of hematopoietic stem cells (HSCs) in the bone marrow (BM) to sense peripheral inflammation and to differentiate preferentially toward the myeloid lineage (Chavakis et al., 2019; King and Goodell, 2011). Moreoever, increased HSC proliferation and skewing towards myelopoiesis have been observed in chronic inflammation such as inflammatory bowel diseases (IBD) and arthritis (Griseri et al., 2012; Oduro et al., 2012). However, the mechanism by which HSCs within the BM are modulated during chronic inflammation, and particularly, the effect of inflammation on the HSC niches are still incompletely understood.

HSCs are maintained by specialized niches in the BM that regulate their fate in terms of quiescence, mobilization, proliferation and differentiation into myeloid and lymphoid lineages. Osteoclasts (OCLs), the bone-resorbing cells and osteoblasts, the bone-forming cells, were shown to contribute to the regulation of HSC niches (Blin-Wakkach et al., 2014; Hanoun and Frenette, 2013). We showed that OCLs promote the formation of HSC niches at birth by controlling the fate of mesenchymal stromal cells (MSCs) that participate in these niches (Mansour et al., 2012a, 2012b). Interestingly, in response to stress signals, OCLs secrete proteolytic enzymes such as Matrix Metallopeptidase 9 (Mmp9) and Cathepsin-K (Ctsk) that are involved in the remodeling of the niches and the mobilization of HSCs outside of their niches (Kollet et al., 2006). Furthermore, in steady state, the in vivo blockage of OCL function reduced HSC numbers in the bone marrow (Lymperi et al., 2011). More recently, an elegant study of dynamic HSC behavior in vivo showed that expansion of HSCs was observed almost exclusively in BM area with bone resorbing activity (Christodoulou et al., 2020). Overall, these observations underline the fundamental role of OCLs in the regulation of HSC niches. Howerver, up to now, the OCL function has been explored only in the context of the on-demand HSC mobilization for emergency hematopoiesis but never in the context of chronic inflammation. However, dysregulated hematopoiesis with myeloid skewing were observed in many chronic inflammatory diseases where OCL activation is increased such as IBD and arthritis. Participation of OCLs to HSC reactivation and myeloid differentiation could therefore be an essential but neglected mechanism for initiating and sustaining chronic inflammation that remains to be elucidated.

Here, to address the contribution of OCLs to HSC activation and myeloid skewing, we used the well characterized model of colitis induced in Rag1−/− mice by transfer of naive CD4^+^ T cells. We showed that OCLs in colitic mice differ from those from controls by a higher expression of genes involved in bone resorption and in particular proteolytic enzymes known to remodel HSC niches. We demonstrated for the first time that OCLs induce hematopoietic progenitor proliferation accompanied by a substantial myeloid skewing in the early phases of colitis. Moreover, we showed that early inhibition of OCLs during chronic colitis dramatically reduce myeloid cell activation and chronic inflammation. Together, these data identify OCLs as potent regulators of HSC homeostasis and may represent a promising target in chronic inflammation.

## Results

### Increased osteoclastogenesis is established before the clinical onset of colitis

To address the participation of OCLs in inflammation *in vivo*, we used the well-characterized model of IBD induced by transfer of naive T cell-transfer into Rag 1−/− mice. In this model, mice develop wasting disease and colitis with high colitis score (7 to 10) associated with splenomegaly, myeloid infiltration and accumulation in the colon, usually in about 6-8 weeks after naive CD4^+^ T cell-transfer (Izcue et al., 2008; Powrie et al., 1996; Wakkach et al., 2008). To better understand the contribution of OCLs to the early phase of intestinal inflammation, we analyzed the relationship between colitis and OCL differentiation during the first weeks after naive CD4^+^ T cell-transfer. As expected, during the first 3 weeks, transferred mice do not develop colitis as indicated by an increased weight (Fig. S1A), the absence of clinical features of colitis and a clinical score below 2 (Fig 1A and Fig. S1B). Interestingly, 3 weeks after naive T cell-transfer, the number of TRAcP^+^ OCLs at the bone surface was already considerably increased leading to decreased trabecular bone in the femur of colitic mice compared to controls (Fig 1B-C). Moreover, among these OCLs, we found an increase of inflammatory-OCLs attested by the expression of Cx3cr1 recently described as marker of these cells (Ibáñez et al., 2016) (Fig 1D). These results indicate that the differentiation of OCLs is established very early and before the clinical onset of colitis in these mice.

**Figure 1.**
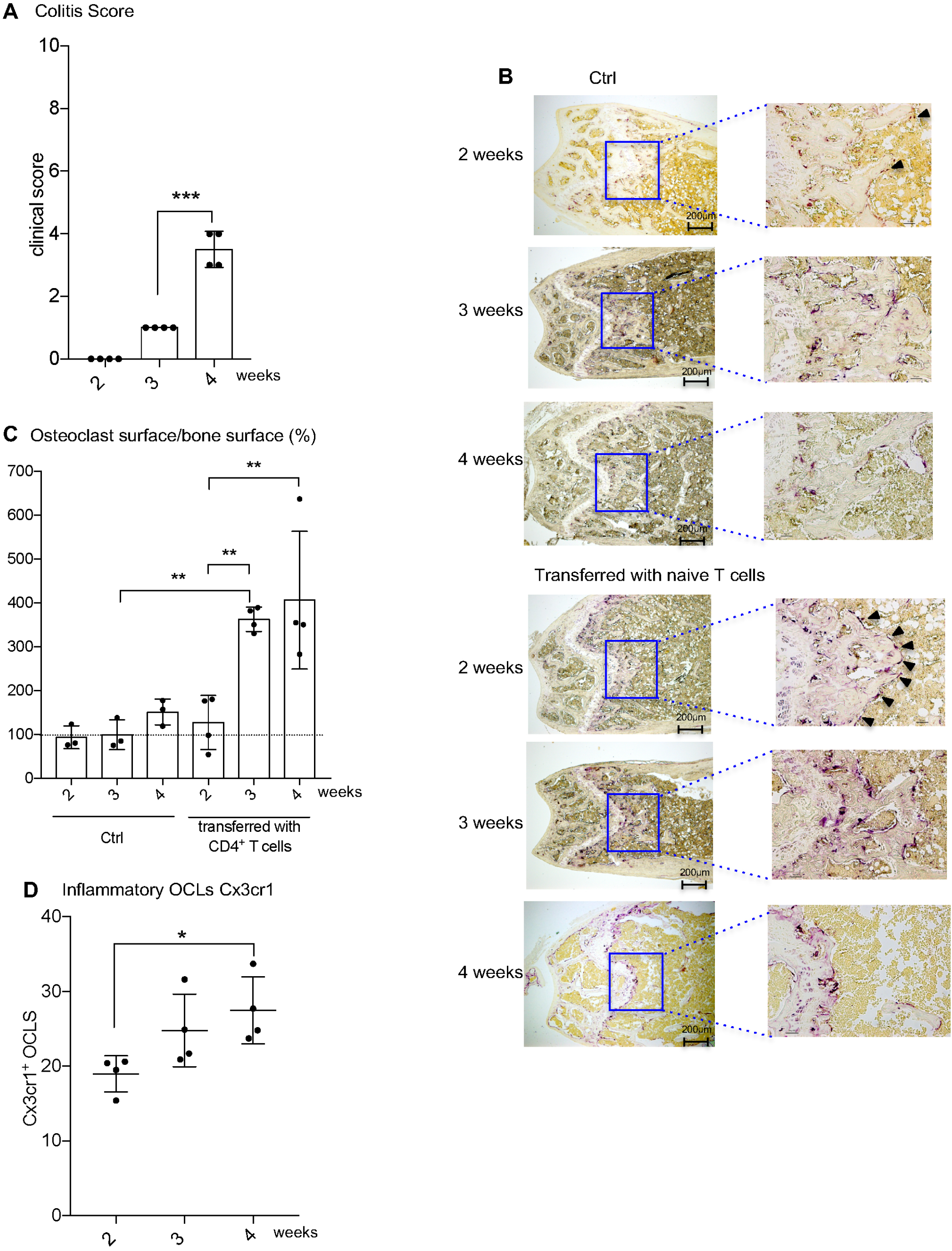
Early OCL differentiation is established before the clinical onset of colitis. Rag1−/− mice received naive CD4 T cells (termed transferred mice) or PBS1X (Ctrl) and were analyzed at the indicated time. (A) Clinical score of transferred *Rag 1−/−* mice. (B) TRAP staining of femora from controls (left) and transferred *Rag 1−/−* mice. Arrowheads indicate TRAP^+^ OCLs. (C) Quantification of TRAP+ OCLs at bone surface. (D) Quantification of FACS analysis of Cx3cr1 expression on OCLs differentiated from BM of transferred *Rag 1−/−* mice at the time indicated in the Figure. Data are representative of at least three independent experiments. *p<0.05; **p<0.01; ***p<0.001

### Comparative transcriptomic analysis revealed that OCLs in colitis are distinct from normal OCLs

To better understand the biological properties of OCLs in colitis on a genome-wide scale, we performed transcriptomic profiling by RNA-seq on sorted OCLs differentiated from the BM of colitic (6 weeks after T cell-transfer) and control mice. A total of 663 genes were significantly differentially expressed (pVal<0.05, Log_2_FC>1) between the 2 populations. Interestingly, Principal Component Analysis (PCA) of the 2000 most expressed genes revealed that colitic and control OCLs were clustered into 2 distinct populations (Fig 2A), which was reinforced by hierarchical clustering analysis (HeatMap) of the top of 57 significantly differentially expressed genes (adj.pVal < 0.05, Log_2_FC > 2) (Fig 2B). These results expand our previous data (Ibáñez et al., 2016) by clearly demonstrating that colitic OCLs differ from control OCLs, confirming thereby the diversity of OCLs.

**Figure 2.**
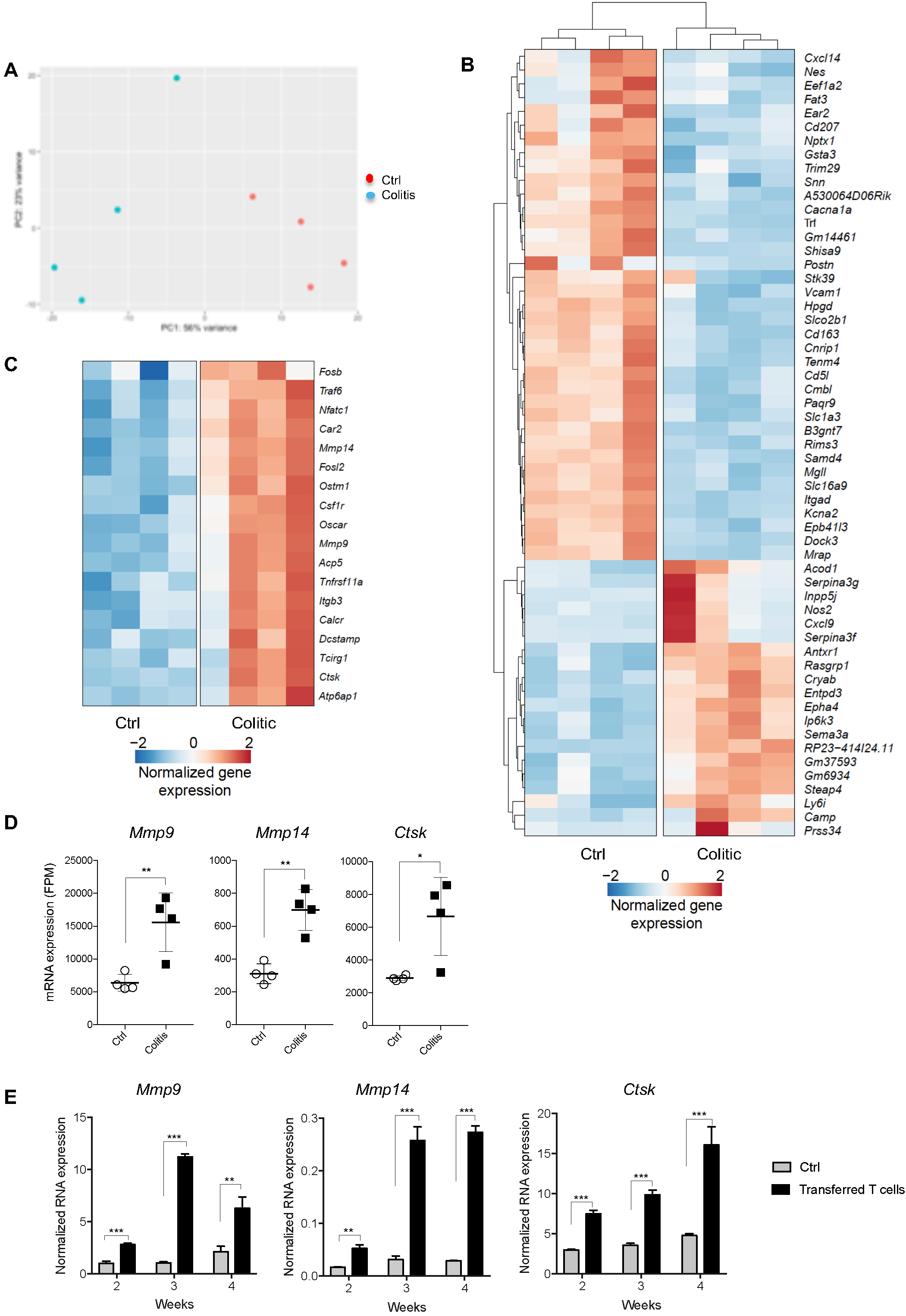
Comparative analysis of purified OCLs from colitic and control mice revealed distinct transcriptomic signatures. Sorted mature OCLs generated from the BM of colitic (6 weeks after T cell transfer) and control mice were compared using RNAseq analysis. (A) Principal component analysis of the genes shows that colitic and control OCLs segregate into 2 distinct clusters. (B) Heatmap visualization of the top 57 differentially expressed genes (pVal<0.05, Log_2_FC>1). (C) Heatmap visualization of selected genes involved in OCL differentiation and bone resorption activity that are significantly increased in colitic OCLs compared to control OCLs (pVal<0.05, Log_2_FC>1). (D) Expression of *Mmp9*, *Mmp14* and *Cstk* in colitic and control OCLs from the RNAseq analysis. Data are expressed as fragments per million (fpm) normalized to total reads per sample. (E) RT-qPCR analysis for the expression of *Mmp9*, *Mmp14* and *Cstk* genes in colitic and control OCLs at week 2, 3 and 4 after transfer of naïve T cells or PBS injection (3 mice for each condition). Results were normalized to the 36B4 gene. *p<0.05; **p<0.01; ***p<0.001.

Furthermore, our transcriptomic profiling revealed that colitic OCLs express higher levels of genes involved in the differentiation and bone resorption function of OCLs (Fig 2C) in line with the increased bone resorption previously observed in colitic mice (Ciucci et al., 2015). Interestingly, among these genes, we observed up-regulation in the expression of cathespsine K (*Ctsk)* and two metallopepetidase genes, *Mmp9* and *Mmp14* (Fig 2D) that have been involved in HSC mobilization and expansion through their capacity to degrade HSC niche components (Kollet et al., 2007; Saw et al., 2019). We then analyzed the expression of these 3 genes by RT-qPCR on purified OCLs derived from the BM of T cell-transferred mice at 2, 3 and 4 weeks. Our results clearly showed a significant increase of the expression of these 3 genes as early as week 2 after the induction of colitis. This increase reached a peak at week 3 with a 10-fold induction for *Mmp9* and *Mmp14* and a 3-fold induction for *Ctsk* (Fig 2E). Collectively, these data demonstrate that modifications in the number and phenotype of OCLs occurred during the first steps of colitis. These results prompted us to analyze whether these OCLs may influence HSCs in terms of expansion and differentiation into myeloid cells, thereby promoting colitis.

### Early OCL activity is associated with HSC expansion and GMP progenitor formation

To address this point, we analyzed the hematopoietic Lin^−^ Sca1^+^ c-kit^+^ (LSK) progenitor cells in the BM of T cell-transferred mice during the first weeks. Interestingly, we observed an increase in the proportion of LSK cells as early as week 2 that continued to grow up to 15-fold at 3 weeks (Fig 3A-B). This accumulation of LSK cells is associated with an increase of their proliferation attested by Ki-67 staining (Fig 3C). These results suggest an alteration of the niche generally linked to LSK cell mobilization, which was confirmed by their increased proportion in the spleen of T cell-transferred mice compared to controls (Fig. S2A-B). Overall, LSK cell proliferation was increased in the BM of T-cell-transferred mice, which was particularly striking at week 3 in parallel with the increased osteoclastogenesis.

**Figure 3.**
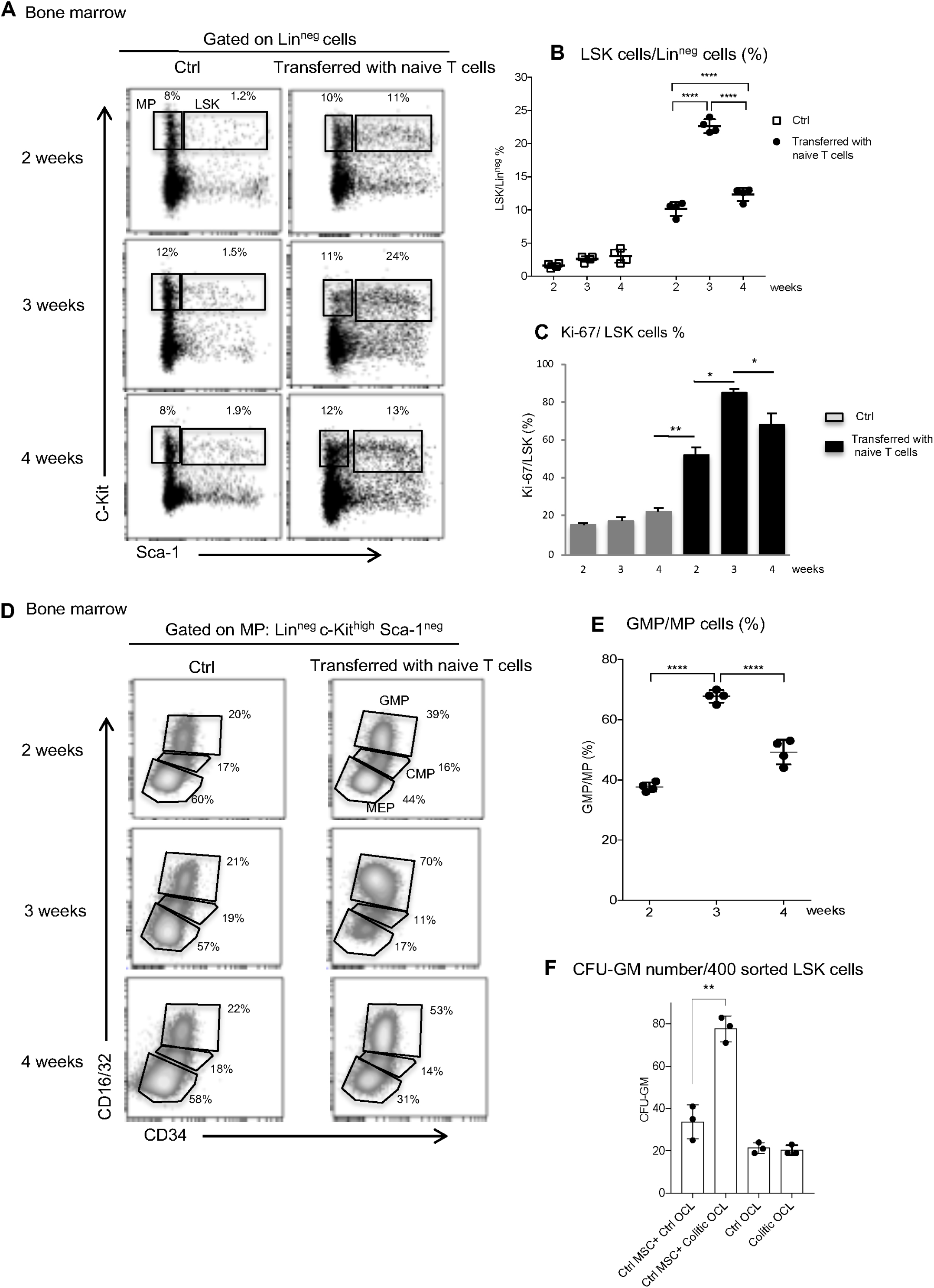
Induction of HSC proliferation and myeloid skewing in the BM of transferred *Rag1−/−* mice. (A) Representative flow cytometry plot and (B) percentage of LSK cells among Lin^−^ cells in BM cells of control (Ctrl) and transferred *Rag1−/−* mice at different time as indicated. (C) BM cells were stained for the proliferation marker Ki-67 and the proportion of Ki-67+ cells among LSK cells was analyzed by flow cytometry. (D) Representative flow cytometry analysis of CMP, GMP and MEP progenitors in the BM. (E) Percentage of GMPs among of MPs. Each point represents an individual mouse. Data are representative of at least three independent experiments. (F) Colony-forming assay (CFU-GM) of sorted LSK cells in co-culture with MSCs from the BM of control mice and in the presence of sorted OCLs differentiated from BM of colitic or control mice. The data are presented as the mean+/-SD of GM-CFU numbers induced by MethoCult GF 3434 medium. Each point represents an individual mouse. Data are representative of two independent experiments. *p<0.05; **p<0.01; ****p<0.0001.

To determine whether the activation of HSC progenitor cells might subsequently induce a change in the orientation of their differentiation, the proportion of common myeloid progenitor (CMP), megacaryocyte-erythrocyte progenitor (MEP) and granulocyte-monocytes progenitors (GMP) were quantified. During the first weeks after T cell transfer in *Rag1*^−/−^ mice, the cell composition within the myeloid progenitor cells (Lin^neg^ c-kit^high^ Sca-1^neg^) changed considerably with a shift to increased GMP compared to control mice that peaked at week 3 (Fig 3D-E). Altogether, the myeloid progenitor composition was dramatically altered with a skew toward GMP differentiation particularly at week 3. The consequence of this change was also observed in the periphery with extramedullary hematopoiesis (Fig. S2A-B) as well as an accumulation of monocytes and neutrophils in the spleen (Fig. S2C-D). We also observed an infiltration of neutrophils in the lamina propria of T cell-transferred mice (Fig. S2E-F).

Hematopoiesis is tightly controlled by the interaction between hematopoietic progenitors and mesenchymal stromal cells (MSCs). To evaluate whether OCLs can modulate the interaction between LSK cells and MSCs, we then performed colony forming assay of sorted LSK cells, in co-culture with MSCs from control mice and OCLs from colitic or control mice. Our results showed that the presence of purified colitic OCLs is sufficient to dramatically enhance the number of CFU-GM from LSK cells (Fig 3F). In the absence of MSCs, OCLs did not affect the number CFU-GM (Fig 3F). Thus, these data demonstrate that colitic OCLs are major modulators of the interaction between LSK cells and their niche and participate in increasing their myeloid differentiation capacity.

### Progenitor expansion and myeloid skewing are induced by colitic OCLs

We next confirmed these results *in vivo* by inducing OCL formation by RANKL injection in normal mice as previously described (Kollet et al., 2006). This approach has been already shown to induce the formation of OCLs (Ibáñez et al., 2016). As expected and already described (Kollet et al., 2006), the number of TRAcP^+^ OCLs on bone surface was elevated in treated mice compared to controls (data not shown). We also observed an increase in the percentage and absolute number of LSK cells (Fig 4A), confirming the data previously reported (Kollet et al., 2006). Interestingly, we also observed a shift in hematopoiesis towards increased GMP formation in mice injected with RANKL (Fig 4B), confirming that increased OCL formation is associated with myeloid skewing *in vivo*.

**Figure 4.**
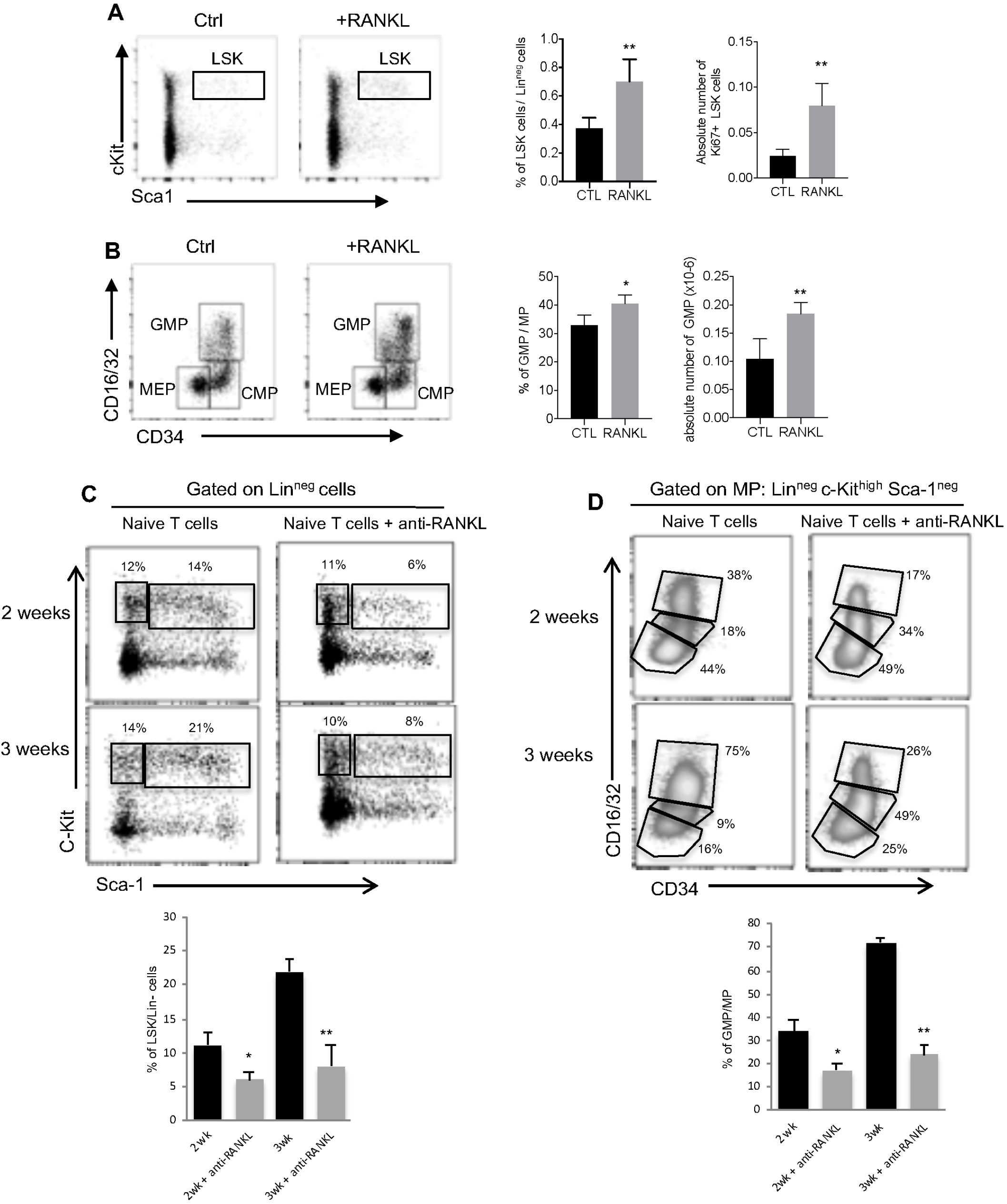
Colitic OCLs promote hematopoietic progenitor mobilization and myeloid skewing. (A-B) Representative flow cytometry analysis (left), percentage (middle) and absolute numbers (right) of LSK cells and GMPs in the BM of wild type mice injected with RANKL. Data are representative of three independent experiments. (C-D) *Rag1−/−* mice were injected 24h after naive CD4 T cell transfer with anti-RANKL twice per week for 2 and 3 weeks. (C) Representative flow cytometry analysis and (D) percentage of LSK cells and GMPs in the BM of untreated and treated transferred *Rag1−/−* mice. Data are representative of two independent experiments. *p<0.05; **p<0.01.

We next blocked RANKL-induced OCL formation in Rag1−/− mice transferred with naive T cells to evaluate the effect of OCLs on hematopoietic progenitor expansion *in vivo*. LSK cell and GMP percentages were significantly decreased in the BM of T cell transferred mice injected with anti-RANKL antibody compared to mice injected with an isotype control antibody (Fig 4C and D).

Collectively, these data show a clear role of colitic OCLs in the proliferation and myeloid differentiation of LSK cells, which could place the modification observed in OCLs as a key downstream colitogenic mechanism leading to increase inflammatory myelopoiesis and colitis.

### Long term blockade of colitic OCLs controls intestinal inflammation and ameliorates bone loss

To assess the contribution of OCL activity in the development of intestinal inflammation, we treated T cell-transferred *Rag1*^−/−^ mice either with zoledronic acid (ZA), a bisphosphonate that inhibits OCL activity (Mansour et al., 2012a; Russell et al., 2008), or with TNF-α inhibitor (TNFi) (Etanercept) used to inhibit intestinal inflammation, 2 weeks after T cell-transfer as indicated in Fig 6A.

As expected, the injection of TNFi inhibited the development of colitis, as revealed by the absence of weight loss (Fig 5A) and low clinical score (Fig 5B). Interestingly, ZA-treated mice maintained a stable weight and were significantly protected from the wasting disease compared to IBD mice, attested by a reduced clinical score compared to colitic mice (Fig 5A-B). Quantification of TRAcP^+^ OCLs on bone section of treated mice in histological analysis of the femur confirmed that ZA as well as TNFi dramatically decreased the number of OCLs on bone surface (Fig 5C). In line with this result, micro-CT analysis revealed that compared to IBD mice, transferred *Rag1*^−/−^ mice treated by both ZA or TNFi display an improvement of bone characterized by a higher BV/TV ratio (Fig 5D). These results strongly suggest that targeting inflammatory OCLs contribute to the control of intestinal inflammation. Accordingly, treatment with two anti-osteoclastogenic agents (ZA and Calcitonine (Calci)) showed a decrease in the percentage of LSK cells accompanied with a decrease in their proliferation (Fig 6A). This result further supports the role of colitis-induced OCLs in the proliferation and myeloid skewing of LSK cells. Of note, long-term blockade of colitis-induced OCLs resulted in a significant decrease in GMP percentage from 60% to 20% and 25% in BM after ZA and Calci treatments, respectively (Fig 6B).

**Figure 5.**
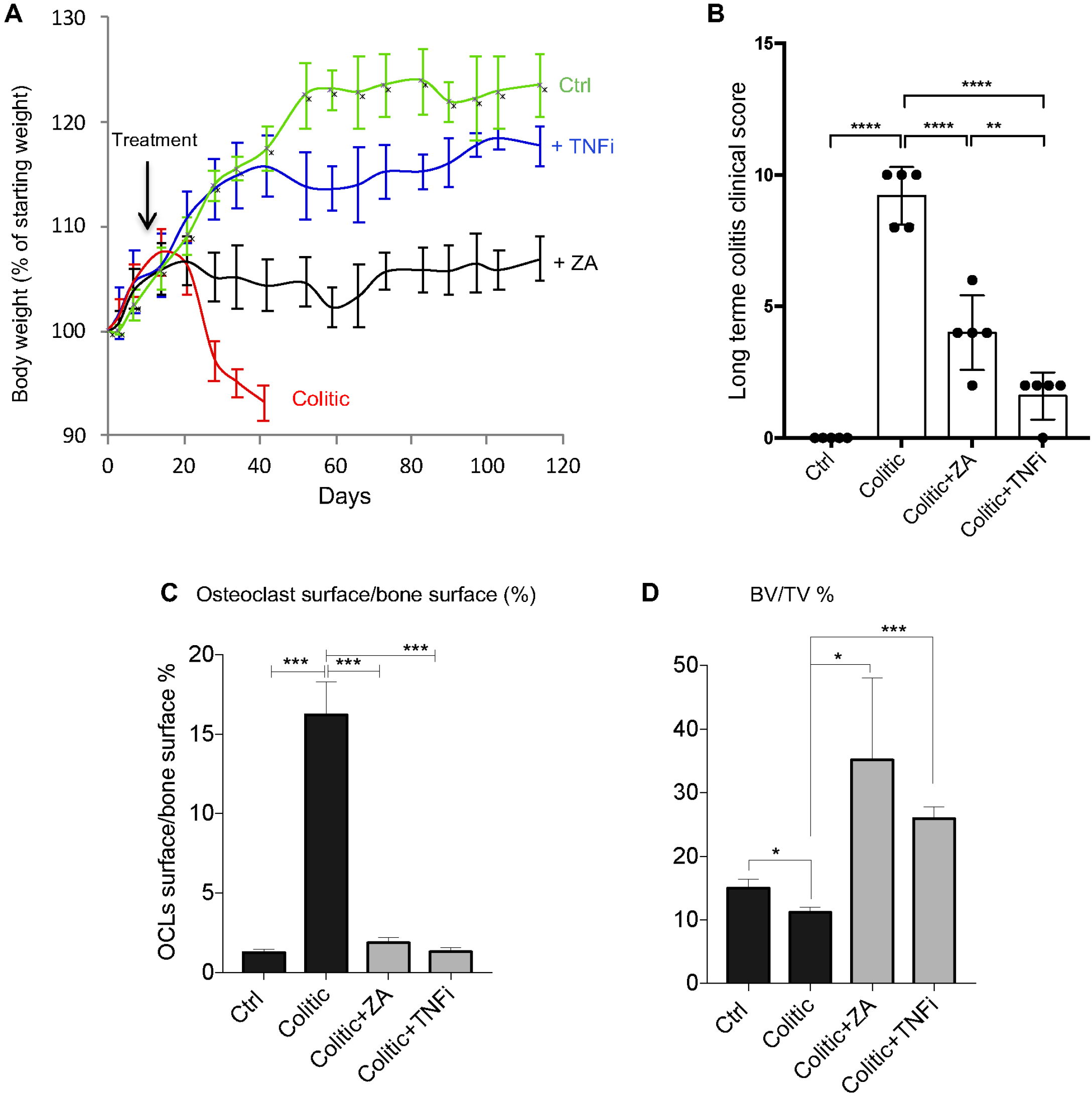
Long term blockade of colitic OCLs controls intestinal inflammation and ameliorate bone loss. (A) Transferred *Rag1−/−* mice were injected twice per week with Etanercept (TNFi), zoledronic acid (ZA), or PBS 1X (colitic), starting from week 2 (arrow “treatment”). The graph represents the mean weight normalized per the initial weight of each mice of 5 animals per group, according to the time. (B) Colitis score 110 days after TNFi and ZA treatment. All results are representative of data generated in at least three independent experiments. (C) Quantification of TRAP+ OCLs at bone surface in the different groups. (D) Percentage of bone volume fraction (BV/TV) of femur analyzed by μCT. All results are representative of data generated in three independent experiments. *p<0.05; **p<0.01; ***p<0.001; ****p<0.0001.

**Figure 6.**
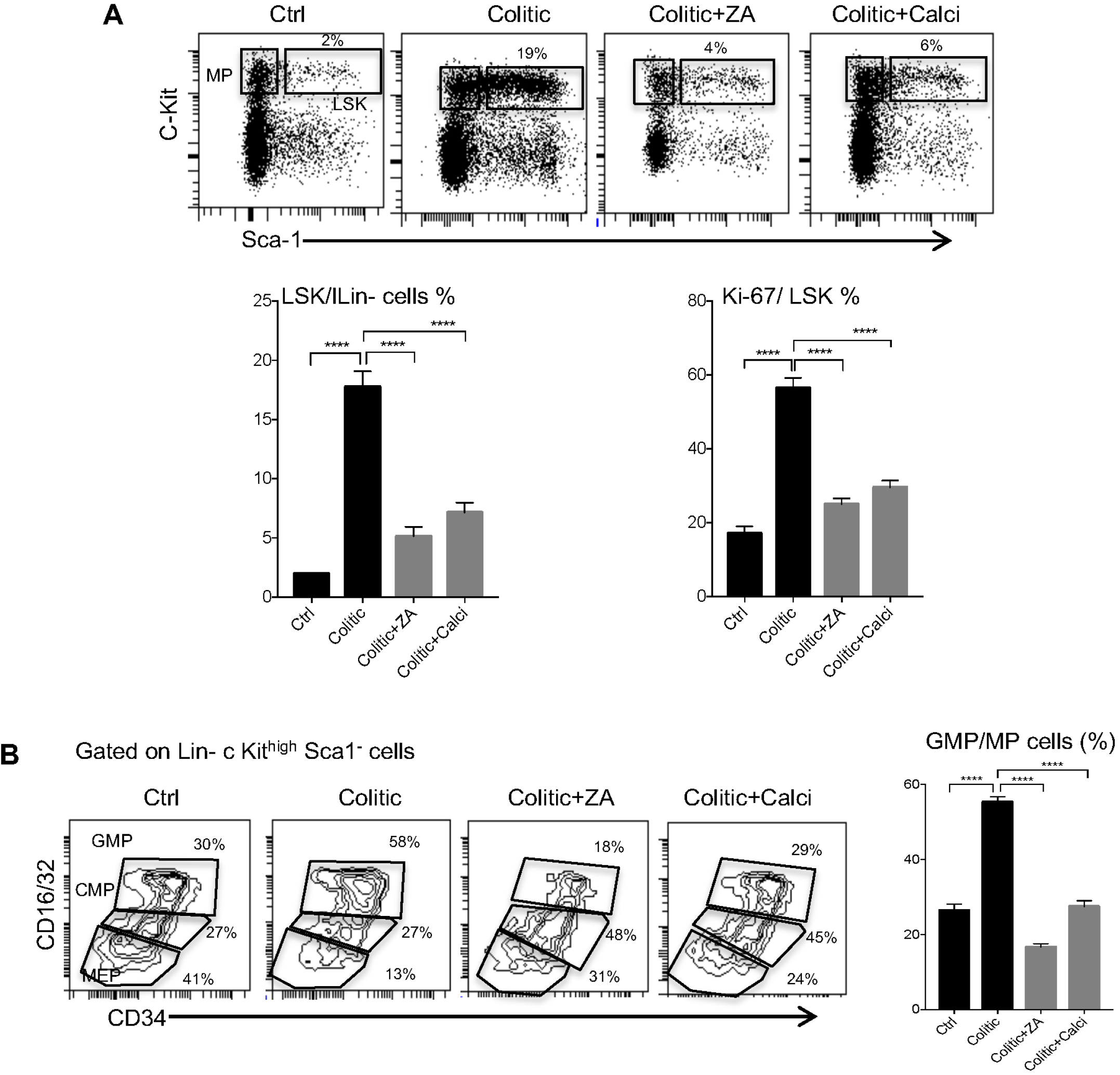
Colitic OCL blockade prevents hematopoietic deregulation and myeloid skewing. Transferred *Rag1−/−* mice were injected twice per week with ZA, Calcitonin (Calci), or PBS 1X (colitic). (A) Representative flow cytometry analysis, percentage and proliferation (Ki-67 staining) of LSKs. (B) Representative flow cytometry analysis of CMP, GMP and MEP progenitors in the BM. The percentage of GMPs among of MPs is indicated in right panel. All results are representative of data generated in three independent experiments. ****p<0.0001.

To further assess the contribution of OCL activity in the development of intestinal inflammation and myeloid differentiation arising from GMPs, we analyzed the myeloid lineage infiltration in the colon of colitis mice and mice treated with ZA and Calci. Colon of colitis mice present a major cell infiltration (Fig S3A), with a marked overabundance of dendritic cells, eosinophils, neutrophils and inflammatory monocytes (Fig S3B-D) as well as an accumulation of colitogenic Th1 cells (Fig. S3E). Interestingly, blockade of colitic OCLs (by ZA and Calci) led to a significant decrease of inflammatory immune cells in the colon of treated mice (dendritic cells, eosinophils, neutrophils, inflammatory monocytes and Th1) (Fig. S3 B-E). Together, our results strongly support a role for colitis-induced OCLs in the onset and exacerbation of intestinal inflammation suggesting that OCLs could represent a new potential therapeutic target in colitis.

### Increased in myeloid dendritic cells and pro-inflammatory monocytes in Crohn’s disease patients correlates with high OCL activity

We next evaluated the link between OCL activity and myelopoiesis in patients affected with Crohn’s disease (CD). Serum concentration of Crosslaps (CTx), the carboxyterminal telopeptide region of type I collagen alpha (1) chain cleaved by OCLs during bone resorption, reflects the osteoclastic activity (Christgau et al., 2000). CTx levels, were measured in the serum of 6 patients with CD (recently diagnosed) and 6 healthy donors, were increased in CD patients compared to controls (Fig 7A). In parallel, we observed a significant increase in the percentage of myeloid BDCA-1^+^ DCs in the blood of CD patients compared to the controls (Fig 7B). In agreement with recent studies (Hanai et al., 2008; Koch et al., 2010), a significant increase of intermediate CD14^+^ CD16^+^ blood monocytes was also observed in patients with CD versus controls (Fig 7C). We next asked whether this increase in inflammatory myeloid cells in CD disease correlated with OCL activity. In contrast to the classical monocytes, there was a strong correlation between CTx and the blood proportion of BDCA-1^+^ DCs (R^2^ = 0.94), as well as intermediate monocytes (R^2^ = 0.78) (Fig 7D). These data suggest that, as in mice, increased OCL activity in CD patients could participate in increased myelopoiesis.

**Figure 7.**
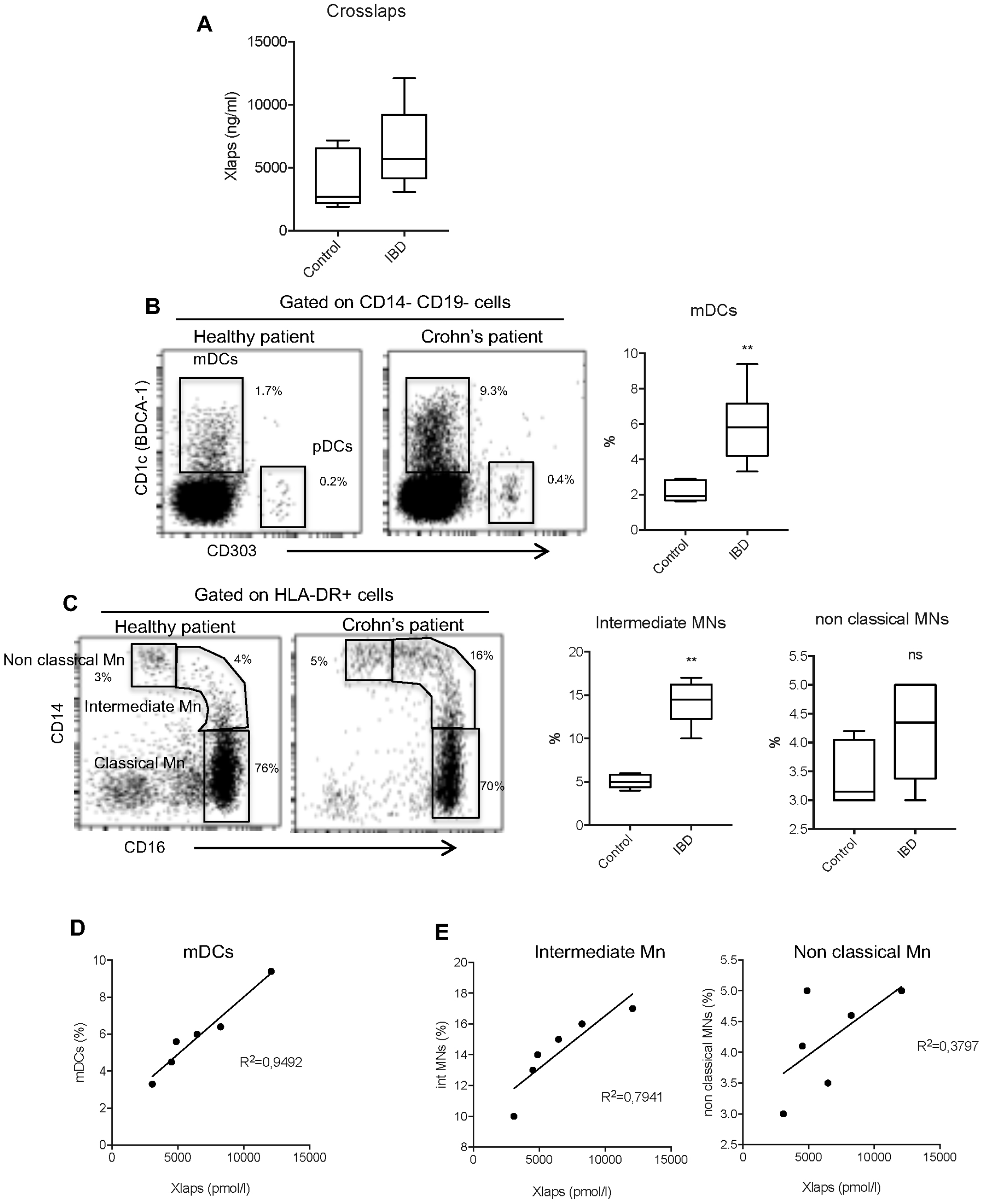
Increase of myeloid dendritic cells and pro-inflammatory monocytes in Crohn’s disease (CD) patients is correlated with OCL activity. (A) Serum concentration level of CTx in healthy donor and patients with CD. Frequency of dendritic cells (DCs) (B) and monocytes (Mn) (C) in the peripheral blood from healthy donor or patients with CD were assessed by flow cytometry. Correlation between CTx and myeloid DCs (D) or monocytes subsets (E) was analyzed using the spearman’s rank correlation test.

## Discussion

The role of hematopoietic progenitors (HSCs) as central determinants of the host response in chronic diseases has long been underappreciated. Here, we identified an unexpected and unique role of osteoclasts in promoting inflammation during the first step of colitis. We showed that OCLs differ between colitis and control conditions. We demonstrated for the first time that colitic OCLs induce HSC proliferation and differentiation markedly towards a substantial myeloid skewing in the early phases of colitis by altering the interactions of HSCs with their niches, which contribute to the disease. Direct inhibition of OCLs by anti-RANKL, calcitonin or bisphosphonate treatment (zoledronic acid) provides strong evidence for the central role of OCLs in controlling HSC proliferation, as well as myeloid skewing in colitis. Interestingly, long term OCL inhibition resulted in decrease of clinical score of colitis, highlighting a beneficial effect on intestinal chronic inflammation. We further document a strong correlation between OCL activity and an increase of inflammatory myeloid cells in the blood of Crohn patients suggesting a role of OCLs in the generation of these pro-inflammatory myeloid cells.

Dysregulation of HSCs with myeloid skewing has been observed in many chronic inflammatory diseases including colitis, rheumatoid arthritis and spondyloarthritis and may represent a shared pathophysiological mechanism (Griseri et al., 2012; Oduro et al., 2012; Regan-Komito et al., 2020). In colitis, inflammatory cytokines control directly distinct components of the aberrant hematopoietic response. For instance IFN-γ increases HSC activity and GM-CSF is involved in the accumulation of GMPs (Griseri et al., 2012). Altogether, our results showed that the dysregulation of HSCs to myelopoiesis occurs before the onset of colitis. Indeed, colitic OCL activity promotes HSC expansion with myeloid skewing before the clinical onset of colitis, which generates a destructive feed-forward loop in which increased numbers of inflammatory myeloid cells enhance inflammatory cytokines, which in turn perpetuates an aberrant hematopoiesis response and an increased osteoclastogenesis leading to osteoporosis.

Recent data highlighted the heterogeneity of OCLs depending on their environment and origin (Ibáñez et al., 2016; Madel et al., 2019, 2020). RNAseq analysis revealed that OCLs from colitic and control mice were clustered into 2 distinct populations providing further evidence of OCL heterogeneity and uncovering a hitherto disregarded OCL diversity that contributes to a deeper understanding of the balance between inflammation and immunosuppression in the BM. Interestingly, transcriptomic profiling of colitic OCLs revealed an overexpression of genes involved in bone resorption in the line with high level of bone destruction observed in colitic mice and IBD patients (Ciucci et al., 2015; Tilg et al., 2008). In particular *Mmp9*, *Mmp14* and *Ctsk*, proteolytic enzymes known to induce mobilization of BM progenitors (Hoggatt and Pelus, 2011; Hoggatt et al., 2018), were increased during the first weeks before the onset of colitis. Mmp9 and Mmp14 expressed in OCLs jointly participate in bone resorption making them key regulators of OCL activity *in vivo* (Zhu et al., 2020). Furthermore, the activity of Mmp9 and Mmp14 disrupt major niche factors, such as Cxcl12 and kit ligand (SCF) involved in HSC retention and homeostasis, which facilitates release and egress of HSCs (Saw et al., 2019). In the light of our observations, the increased expression by colitic OCLs of the Mmp9 / Mmp14 couple may represent a molecular mechanism by which the colitic OCLs induce the expansion of HSCs during the onset of colitis. In other hand, Ctsk, the key marker of bone degradation, also has an important role in regulation of niche factors such as Cxcl12 and in progenitor mobilization, together with Mmp9, in stress condition (Kollet et al., 2006). Overall, our results suggested that overexpression of Ctsk and Mmp9/Mmp14 in colitic OCLs, each with a variety of overlapping substrates involved in HSC niches, may contribute to a powerful regulation of the niche and thus facilitate HSC mobilization and proliferation observed during first weeks before onset of colitis.

Mesenchymal cells (MSCs) are key components of the HSC niche. We previously showed that OCL activity can modulate MSC function and control their fate at birth for establishing BM HSC niches *in vivo* (Mansour et al., 2012a). Our *in vitro* studies indicated that BM-MSCs support and increased the differentiation of LSKs into GMP solely in the presence of colitic OCLs attested by CFU-GM assay. These results demonstrated that colitic OCLs alter the phenotype of BM-MSCs allowing them to orientate hematopoietic differentiation towards myelopoiesis, but the precise mechanism involved remain to be determined.

Actually, anti-resorptive treatments such as bisphosphonates are used to treat bone resorption due to IBD in patients (Hu et al., 2017). As bone resorption is coupled with bone formation, these long-term treatments also inhibit bone remodeling and dramatically affect the bone quality, increasing the risk of fracture and osteonecrosis. Our study highlighted the heterogeneity of OCLs and particularly their involvement in colitis. We believe that targeting only the colitic OCL subset, in the first steps of IBD, will avoid the complications of actual anti-resorptive treatments that block the activity of all OCLs and will ameliorate IBD.

Collectively, our results shed light on a unique role of OCLs in colitis and suggest that they may exert their colitogenic activity by acting on the initial step of hematopoiesis by increasing myelopoiesis. Altogether, targeting OCLs responsible for the augmentation of inflammatory myeloid cells may represent a promising approach for limiting inflammation and complementary therapeutic strategy for the early treatment of IBD. Ultimately, we believe that precise characterization of OCLs that control the function of hematopoietic system will offer opportunities to develop novel therapeutics in IBD but also in other chronic inflammatory diseases with a strong myeloid component such as rheumatoid arthritis and spondylarthritis. It will also improve our general understanding of the regulation of hematopoiesis.

## Methods

### Mice

C57BL/6J mice were purchased from Harlan Laboratory (France), and C57BL/6 Rag1^−/−^ mice from Charles River Laboratory (France). Animals were maintained in our animal facility in accordance with the general guidelines of the institute, and approval for their use in this study was obtained from the Institutional Ethic Committee for Laboratory Animals (CIEPAL-Azur). Inflammatory osteoclastogenesis was mimicked by i.p. injection of RANKL (R&D) in C57BL/6J mice, at 2μg/injection, twice a day for 3 consecutive days and mice were sacrificed at day 5, as described (Ibáñez et al., 2016; Kollet et al., 2006).

### Induction of colitis

Colitis was induced in Rag1^−/−^ mice by one i.p. injection of 4.10^5^ sorted splenic naive CD4^+^CD25^neg^CD45RB^high^ T cells from C57BL/6J mice as described (Wakkach et al., 2008). Mice were weighed twice a week and any mice losing > 20% of their starting body weight were euthanized. When indicated, mice were i.p injected twice a week with 0.25mg of anti-RANKL (IK22-5) or isotype control (rat IgG2a, eBioscience) or 100 μg/kg Zoledronic Acid (ZA) (Novartis) or 0.25 mg of Etanercept (Embrel) or 20U/kg of calcitonin (Calci)(Novartis). Clinical score was graded semi-quantitatively from 0 to 4 for each of the four following criteria: hunching, weight loss, diarrhea, and colon length as described (Ciucci et al., 2015).

### High resolution μCT

Fixed femora were scanned with high resolution μCT (Viva CT40, Scanco) (EcellFrance facility, Université Montpellier, France) as described in (Ciucci et al., 2015). Data were acquired at 55 keV with a resolution of 10 μm cubic.

### Histological analysis

Femora were fixed in 4% paraformaldehyde for 24h at 4°C, decalcified for 10 days in 10% EDTA at 4°C and incubated overnight in 30% glucose solution. 7-μm bone sections were stained for TRAcP activity (leukocyte acid phosphatase kit, Sigma-Aldrich) according to the manufacturer’s protocol. After hematoxylin staining, images were acquired on a light microscope (Carl Zeiss, Inc.).

### Primary osteoclast culture

Osteoclasts were differentiated from murine BM cells as described previously (Madel et al., 2020), collected from the culture plates using Accutase (Sigma-Aldrich) and used for subsequent FACS analysis and sorting as previously described (Madel et al., 2018).

### RNA-sequencing on sorted osteoclasts

Total RNA was extracted with the RNeasy kit (Qiagen) and 100 ng RNA was processed for directional library preparation (Truseq stranded total RNA library kit, Illumina). Libraries were pooled and sequenced paired-ended for 2×75 cycles on a Nextseq500 sequencer (Illumina) to generate 30-40 million fragments per sample. Total RNA (100 ng) from 4 biological replicates (from 4 different mice) in each group was extracted from *in vitro* differentiated OCLs (after sorting according to their multinucleation as described in (Madel et al., 2018) with the RNeasy kit (Qiagen) and processed for directional library preparation (Truseq stranded total RNA library kit, Illumina). Libraries were pooled and sequenced paired-ended for 2 x 75 cycles on a Nextseq500 sequencer (Illumina) to generate 30–40 million fragments per sample. After quality controls, data analysis was performed with 2 different approaches. For the first one, reads were ‘quasi’ mapped on the reference mouse transcriptome (Gencode vM15) and quantified using the SALMON software with the mapping mode and standard settings (Patro et al., 2017). Estimates of transcripts counts and their confidence intervals were computed using 1000 bootstraps to assess technical variance. Gene expression levels were computed by aggregating the transcript counts for each gene. Gene expression in biological replicates was then compared using a linear model as implemented in Sleuth (Pimentel et al., 2017) and a false discovery rate of 0.01. Lists of differentially expressed genes were annotated using Innate-DB and EnrichR web portals. For the second approach, raw RNAseq fastq reads were trimmed with Trimmomatic and aligned to the reference mouse transcriptome (Gencode mm10) using STAR (v. 2.6.1 c) (Dobin et al., 2013) on the National Institutes of Health high-performance computing Biowulf cluster. Gene-assignment and estimates counts of RNA reads were performed with HTseq (Anders et al., 2015). Further analyses were performed with R software and gene expression in biological replicates (n = 4) was compared between the different conditions to identify differentially expressed genes using DESeq2 (Love et al., 2014) with the Wald test (FDR < 0.01). Gene ontology (GO) pathway analyzes were performed using Goseq (Young et al., 2010). Both approaches gave equivalent results (not shown).

### Gene expression analysis

Total RNA of sorted OCLs differentiated *in vitro* from BM of control and T cell-transferred *Rag1−/−* mice were extracted using TRIzol reagent following manufacturer’s instructions. RT-qPCR was performed after reverse transcription (Superscript II, Life Technologies) as described previously using SYBR Green. List of primers: Mmp9 (5′-TGAGTCCGGCAGACAATCCT-3′; 5′-CGCCCTGGATCTCAGCAATA-3′), Mmp14 (5′-CCC TAG GCC TGG AAC ATT C-3′; 5′-TCT TTG TGG GTG ACC CTG ACT-3′), Ctsk (5′-CAGCAGAGGTGTGTACTATG-3′; 5′-GCGTTGTTCTTATTCCGAGC-3′), Samples of 3 biological replicates were run in triplicates and results were normalized to the reference gene 36B4 (5’-TCCAGGCTTTGGGCATCA-3’; 5’-CTTTATCAGCTGCACATCACTCAGA-3’) using the 2−∆Ct method as described (Mansour et al., 2012a).

### Leukocyte Isolation, antibodies and flow cytometry

Single cell suspensions were prepared from spleen, Lamina Propria (LP) and BM as previously described (Ciucci et al., 2015) and were stained with different antibodies used for FACS (FACS canto, BD Biosciences) are indicated in the figures.

HSCs were analyzed after exclusion of lineage positive (Lin) cells using FITC-conjugated antibodies against CD3, B220, CD11b, Gr-1 and Ter-119, and after staining with PE-Cy7 Sca-1 and PE-c-kit. HSCs were indicated as Lin^−^ Sca-1^+^ c-kit^+^ (LSK). Common myeloid progenitor (CMP), megacaryocyte-erythrocyte progenitor (MEP) and granulocyte-monocytes progenitors (GMP) were analyzed using Lin^−^ Sca-1^−^ c-kit^high^, CD34 and CD16/CD32. For LSK proliferation, cells were fixed and stained with Ki-67.

Antibodies anti-CD3 (17A2), anti-CD4 (H129.19), anti-CD11b (M1/70), CD11c (HL3), CD16/CD32 (2.4G2), CD25 (PC61) CD34 (RAM34), c-kit (2B8), anti-Ly6C (AL-21), anti-B220 (RA3-6B2), anti-Gr1 (RB6-8C5), anti-Ter119 (TER-119) were purchased from BD Biosciences (Le Pont De Claix, France). Purified anti-RANKL (IK22-5) was gift from CG. Mueller (CNRS UPR 3572, Strasbourg, France). Antibodies anti-CD45RB (C363,16A), CD170 (1RNM44N), anti-Cx3cr1 (2A9-1) Sca-1 (D7), and Ki-67 (SOLA 15) were purchased from Thermo Fisher Scientific).

### Colony Forming Assay

Sorted LSK cells from BM of control *Rag 1−/−* mice were suspended in methylcellulose medium and co-cultured with sorted OCLs from BM of colitic or control *Rag 1−/−* mice and with or without control mesenchymal stromal cells (MSCs) to quantify clonogenic colony forming units in triplicate cultures (MethoCult GF M3434, Stem Cell Technologies). Colonies were scored after 10–14 days. CFU-G (granulocyte), M (macrophage), or GM (granulocyte-macrophage) colonies are referred to as CFU-GM.

### Patients

PBMC and serum were isolated from freshly collected, heparinized peripheral blood of healthy voluntary donors and patients with Crohn’s disease (Department of gastroenterology, CHU of Nancy, France). The protocol was approved by the local ethics committee of the Hospital Center of Nancy University and all patients and control subjects who were included in this study included in the study provided a signed informed consent. For the blood staining, mouse anti-human CD14, CD16, CD19, HLA-DR and CD66b were purchased from BD Biosciences for monocytes staining and the blood dendritic cell (DC) enumeration kit was purchased from Miltenyi Biotech.

### Statistical analysis

All data were analyzed using Graph Pad Prism 8.0 software using an appropriate two tailed student’s t-test with Bonferroni adjustment when comparing two groups. When more than two groups were compared two-way analysis of variance (ANOVA) was used. Statistical significance was considered at p<0.05. Experimental values are presented as mean ± standard deviation (SD) of at least three biological replicas. Error bars for gene expression analysis of humans and mice using RT-qPCR show the mean value with 95% confidence interval. All experiments were repeated with a minimum of three biological replicates and at least two technical replicates.

## Acknowledgements

We acknowledge the Genomic Facility of the UFR Simone Veil, (Universite’ Versailles-Saint-Quentin, France) for the RNA sequencing, the IRCAN animal core facility (Nice, France) and the preclinical platform of ECELLFRANCE for microCT analysis (IRMB, Montpellier, France). This work utilized the computational resources of the NIH-HPC-Biowulf cluster (http://hpc.nih.gov). The work was supported by the Agence Nationale de la Recherche (ANR-16-CE14-0030), (ANR-19-CE14-0021-01), the French Association François Aupetit, and the French government, managed by the ANR as part of the Investissement d’Avenir UCA^JEDI^ project (ANR-15-IDEX-01). M-B M is supported by the Fondation pour la Recherche Médicale (FRM, ECO20160736019).

## Author contributions

A.W and C.B-W conceived the project and designed the experiments. M-B.M. L.I and HJ performed experiments, analyzed and interpreted data. T.C and H-J.G. analyzed the RNAseq data. MT performed animal experimentation. L.PB and D.M managed experiments on patients. M.R, C-G.M, L.PB and D.M actively contributed to the discussion of results and provided helpful advices throughout the study.

## Supplemental figure legends

**Fig. S1:**
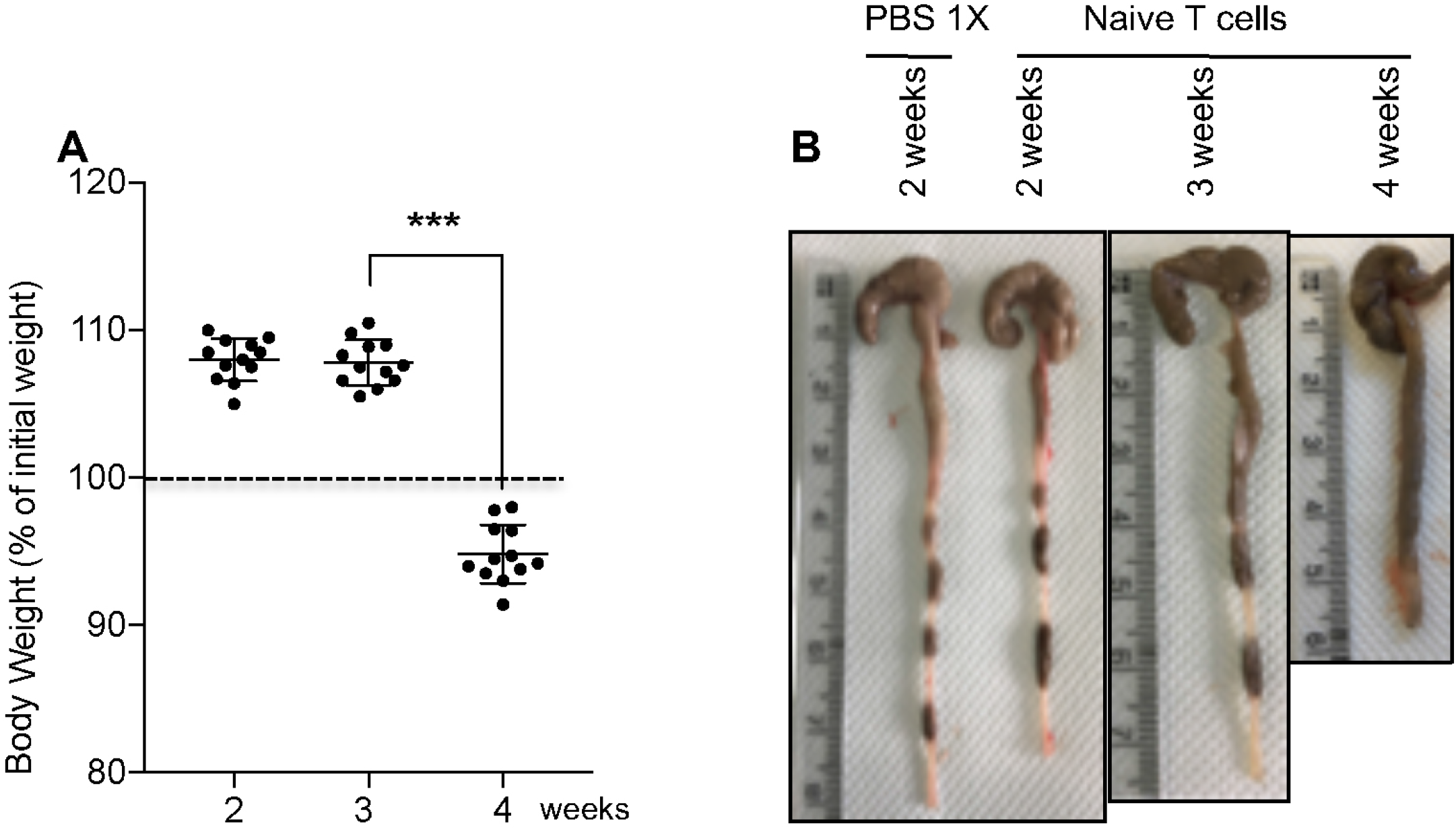
Follow up of Rag1−/− mice in the first weeks after induction of colitis. Rag1−/− mice that were transferred with naive CD4 T cells were monitored during the first weeks after the transfer for (A) changes in their body weight and (B) aspect and length of the colon compared to mice receiving just PBS 1X (representative images).

**Fig. S2:**
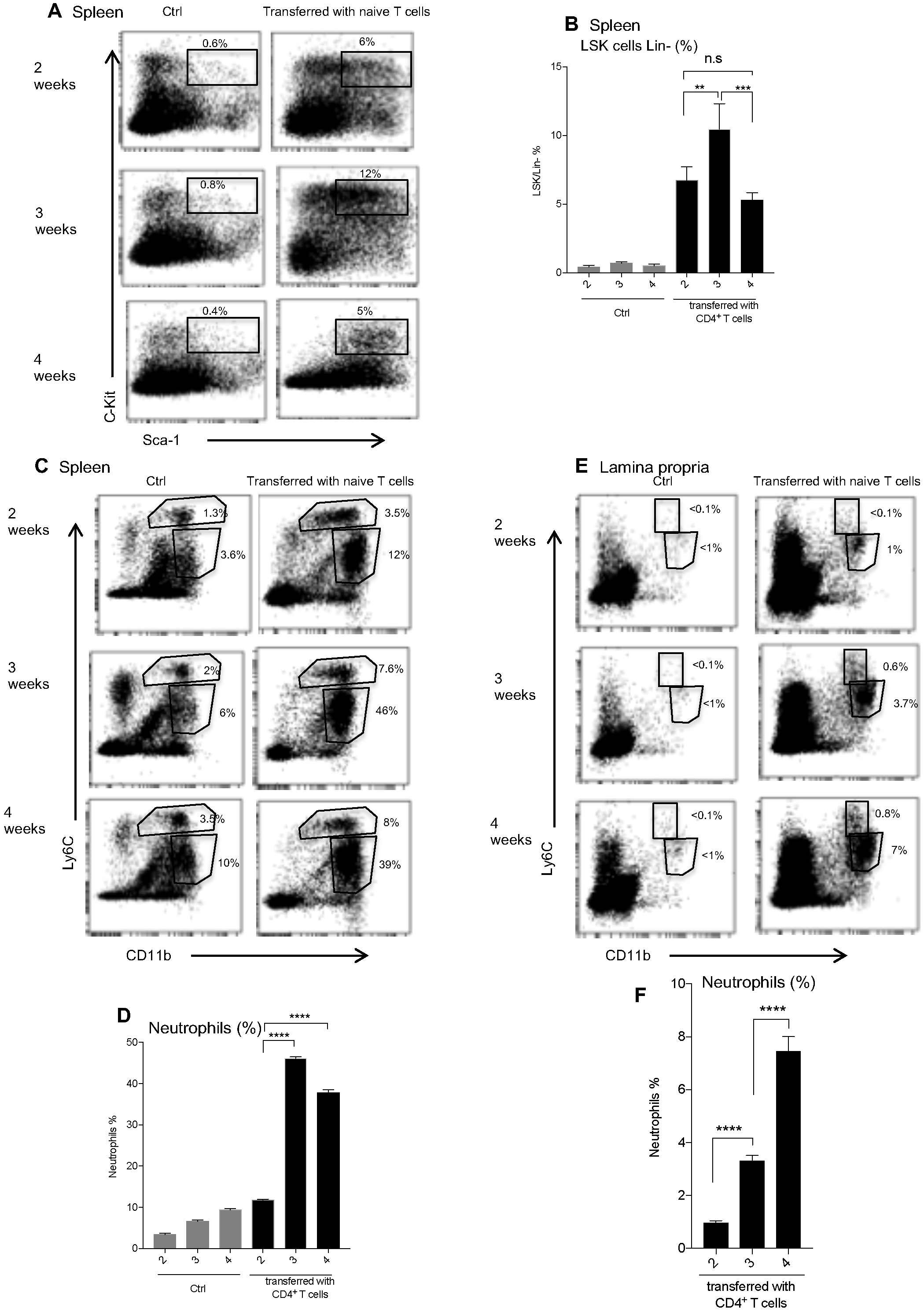
Follow up of Rag1−/− mice in the first weeks after induction of colitis. (A) Representative flow cytometry plot and (B) quantification of the proportion of LSK cells among Lin^−^ cells in the spleen of control (Ctrl) and transferred *Rag1−/−* mice at the indicated time. (C) Representative flow cytometry plot of splenic cells after staining with CD11b and Ly6C antibodies in the spleen of control (Ctrl) and transferred *Rag1−/−* mice at the indicated time. (D) Quantification of the neutrophils (CD11b^+^ Ly6C^low^) in the spleen of control (Ctrl) and transferred *Rag1−/−* mice at the indicated time. (E) FACS analysis of cells from the lamina propria of control (Ctrl) and transferred *Rag1−/−* mice at the indicated time. (F) Quantification of the neutrophils (CD11b^+^ Ly6C^low^) in the lamina propria of control (Ctrl) and transferred *Rag1−/−* mice at the indicated time. Results are representative of the mean ± SD of at least 4 mice in each group. **p<0.01; ***p<0.001; ****p<0.0001.

**Fig. S3:**
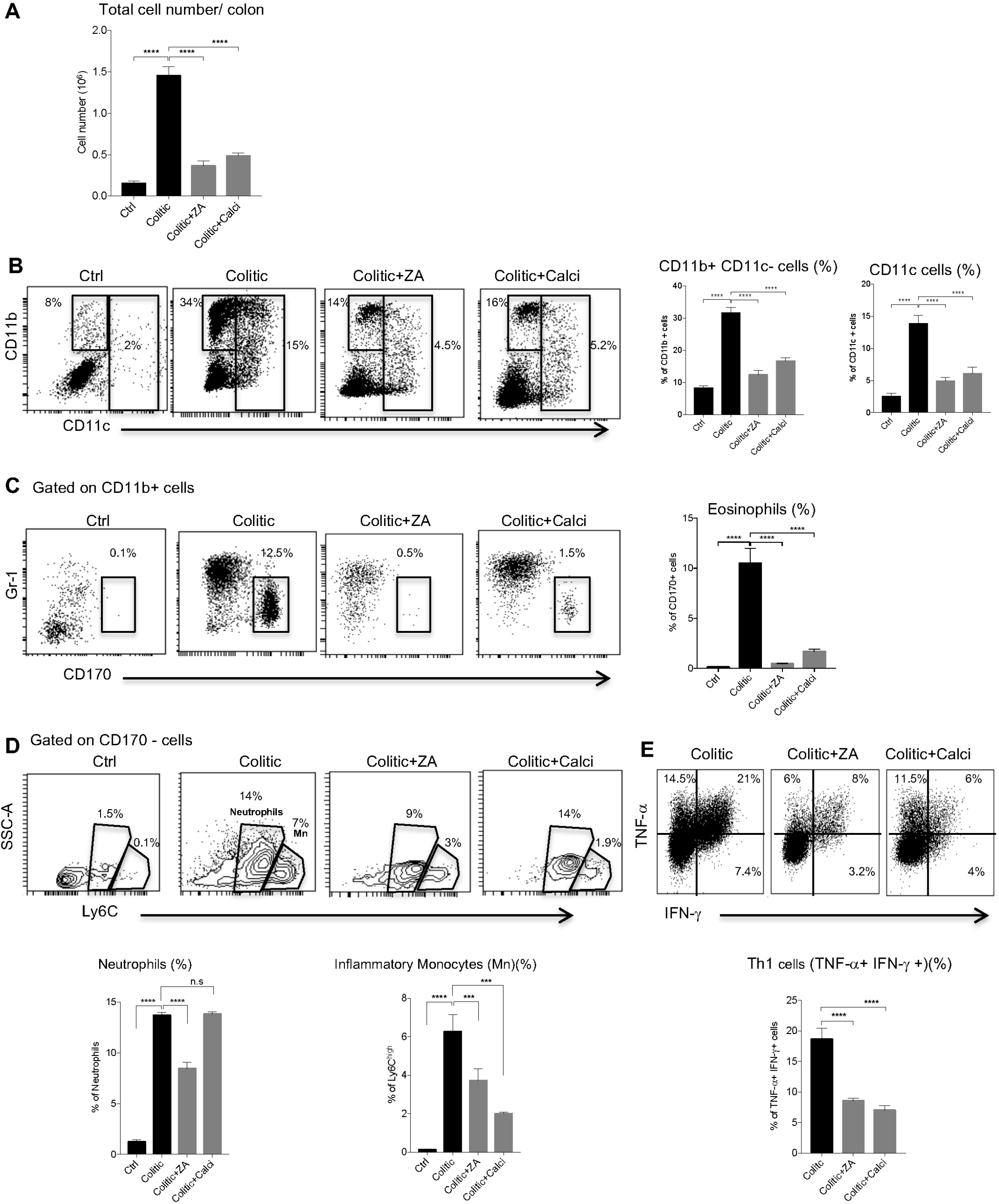
Long-term blockade of colitic OCL limited inflammatory cells recruitment and protected the gut from inflammation. Transferred *Rag1−/−* mice were injected twice per a week with ZA, Calcitonin (Calci) up to 15 weeks, or PBS 1X (colitic). (A) Total cell number in lamina propria. (B) Representative flow cytometry staining of dendritic and myeloid cells with their percentage indicated (right panel). (C) Representative flow cytometry staining of eosinophils and the percentage among CD11b+ cells (right panel). (D) Representative flow cytometry staining of neutrophils and inflammatory monocytes and the percentage among CD11b+ CD170-cells is indicated in the panel at bottom. (E) Representative analysis of intracytoplasmic TNF-α and IFN-γ cytokines of CD4+ T cells in lamina propria and the percentage of TNF-α+ IFN-γ + cells is indicated in the panel at bottom. All results are representative of data generated in three independent experiments. ***p<0.001; ****p<0.0001.

